# Small Animal Jet Injection Technique Results in Enhanced Immunogenicity of Hantavirus DNA Vaccines

**DOI:** 10.1101/2020.11.09.374439

**Authors:** Rebecca L. Brocato, Steven A. Kwilas, Matthew D. Josleyn, Simon Long, Xiankun Zeng, Casey C. Perley, Lucia M. Principe, Brandon Somerville, Melanie V. Cohen, Jay W. Hooper

## Abstract

DNA vaccine evaluation in small animals is hampered by low immunogenicity when the vaccines are delivered using a needle and syringe. To overcome this technical hurdle we tested the possibility that a device developed for human intradermal medicine delivery might be adapted to successfully deliver a DNA vaccine to small animals. The PharmaJet^®^ Tropis device is a FDA 510(k)-cleared disposable syringe, needle-free jet injection device designed to administer medicines to the human dermis in a 0.1 mL volume. Here, we found that hantavirus DNA vaccines administered to Syrian hamsters using Tropis were substantially more immunogenic than the same vaccines delivered by needle/syringe or particle mediated epidermal delivery (gene gun) vaccination. By adjusting how the device was used we could deliver vaccine to either subcutaneous tissues, or through the skin into the muscle. RNA and/or antigen expression was detected in epidermal, subepidermal and fibroblast cells. We directly compared six optimized and non-optimized hantavirus DNA vaccines in hamsters. Optimization, including codon-usage and mRNA stability, did not necessarily result in increased immunogenicity for all vaccines tested; however, optimization of the Andes virus (ANDV) DNA vaccine protected vaccinated hamsters from lethal disease. This is the first time active vaccination with an ANDV DNA vaccine has shown protective efficacy in the hamster model. The adaptation of a human intradermal jet injection device for use as a method of subcutaneous and intramuscular jet injection of DNA vaccines will advance the development of nucleic acid based medical countermeasures for diseases modeled in hamsters.

## Introduction

Hantaviruses cause two diseases, hantavirus pulmonary syndrome (HPS), also known as hantavirus cardiopulmonary syndrome, and hemorrhagic fever with renal syndrome (HFRS). The etiological agents of HPS are primarily New World hantaviruses such as Andes virus (ANDV) and Sin Nombre virus (SNV) [1]. These viruses target the lungs resulting in a disease characterized by fever, vascular leakage, tachypnea, shock, pulmonary edema, and cardiac failure [2, 3] with an associated case fatality rate of 35 to 40% [4, 5]. The etiological agents of HFRS are Old World hantaviruses such as Hantaan virus (HTNV), Puumala virus (PUUV), Dobrava virus (DOBV) and Seoul virus (SEOV) [1]. These viruses target the kidney and have lower case fatality rates, up to 15% for HTNV- and <1% for PUUV-associated HFRS, respectively [6]. There are currently no FDA-approved vaccines for the prevention of hantavirus diseases [7].

Hantaviruses are tri-segmented, negative sense, RNA viruses. The three genome segments are denoted as small (S), medium (M), and large (L), and encode the nucleocapsid, the glycoproteins Gn and Gc, and the RNA-dependent RNA polymerase, respectively [1]. Our laboratory has generated DNA vaccines targeting the M segment of ANDV [8], SNV [9], HTNV [10], PUUV [11], DOBV [12] and SEOV [13]. Early studies testing the immunogenicity of these vaccines using particle-mediated epidermal delivery (gene gun) revealed that SNV, HTNV, PUUV, and SEOV DNA vaccines were immunogenic in hamsters, and the ANDV DNA vaccine was not [8-11]. In addition, when the HTNV and ANDV DNA vaccines were combined into a single plasmid, this resulted in a dominant negative effect where hamsters did not develop an antibody response to either HTNV or ANDV [14]. All of these vaccines, including the ANDV DNA vaccine, have demonstrated immunogenicity in other species, namely rabbits and nonhuman primates [8-11, 14, 15]. The HTNV and PUUV DNA vaccines have been evaluated in phase 1 trials using gene gun [16], or intramuscular (IM) electroporation [17]. A combination HTNV/PUUV DNA vaccine delivered by IM electroporation is in an ongoing clinical trial (NTC03718130). Phase 1 trials of the HTNV and PUUV DNA vaccines (S-15-40 2289) and an ANDV DNA vaccine (NTC03682107) administered using jet injection are in progress [18].

Pre-clinical vaccine research can be greatly accelerated and de-risked using rodent models. Immunogenicity and efficacy data can be generated if appropriate animal models exist. For some diseases there are no large animal models but small animal models do exist. For example, ANDV infection in hamsters, but not nonhuman primates or any other tested species, results in a lethal disease that recapitulates human HPS [19]. Previously, vaccination of small animals such as mice or hamsters were limited by volume of the inoculum and the capacity of the vaccination device. The gene guns, both ND10 and XR1 particle-mediated epidermal delivery device, fit this criteria using plasmid DNA precipitated onto gold beads [8, 11]. Compressed helium was used for vaccine delivery. More recently, we have used disposable needle-free jet injection (PharmaJet^®^) devices to evaluate hantavirus vaccines in nonhuman primates and rabbits. Delivery of ANDV and SNV DNA vaccines with the PharmaJet^®^ Stratis device elicited statistically significant increases in antibody titers when compared to IM needle/syringe delivery [15]. Unfortunately, delivery of 0.5 mL volume makes this device prohibitive in small animals. The PharmaJet^®^ Tropis device delivers 0.1 mL by intradermal injection in humans. We reasoned that it could be possible that this jet injection device could be adapted to deliver DNA to a small animal more efficiently than needle and syringe. We found that two different techniques could be used to effectively deliver multiple hantavirus DNA vaccines to Syrian hamsters.

## Materials and Methods

### Viruses, cells, and medium

ANDV strain Chile-9717869 [20], SNV strain CC107 [21], SEOV strain SR-11 [22], HTNV strain 76-118, PUUV strain K27, and DOBV strain Dobrava [10] were propagated in Vero E6 cells (Vero C1008; ATCC CRL 1586). The VSVΔG*rLuc pseudovirion (PsV) is a recombinant VSV derived from a full-length cDNA clone of the VSV Indiana serotype in which the G-protein gene has been replaced with the Renilla luciferase gene. ANDV, SNV, SEOV, HTNV, PUUV, and DOBV PsV were produced in human embryonic kidney (HEK) 293T cells using DNA vaccine plasmids pWRG/AND-M(opt2), pWRG/SN-M(opt), pWRG/SEO-M(opt2), pWRG/HTN-M(co), pWRG/PUU-M(s2), and pWRG/DOB-M(opt), respectively, by methods previously described [23]. Vero, Vero E6, and HEK 293T cells were maintained in Eagle’s minimal essential medium with Earle’s salts (EMEM) containing 10% fetal bovine serum (FBS), 10mM HEPES pH 7.4, and antibiotics (penicillin [100 U/mL], streptomycin [100 μg/mL]) (cEMEM) at 37°C in a 50% CO2 incubator.

### Hantavirus vaccines and methodology

ANDV, SNV, SEOV, HTNV, PUUV, and DOBV DNA vaccine plasmids used were pWRG/AND-M(1.1), pWRG/AND-M(opt2), pWRG/SN-M(opt), pWRG/SEO-M(opt2), pWRG/HTN-M(x), pWRG/HTN-M(co), pWRG/PUU-M(s2), and pWRG/DOB-M(opt). The PharmaJet^®^ Tropis device was used for both subcutaneous (SC) and IM vaccinations in hamsters. For the SC vaccination, hamsters were anesthetized by inhalation of vaporized isoflurane using an IMPAC 6 veterinary anesthesia machine. Once anesthetized, fur on the abdomen was removing using a pair of clippers. The abdominal skin was folded over itself and held in place using a double-gloved thumb and forefinger. The PharmaJet^®^ Tropis was placed on the skin resting over the forefinger and the device was activated. For the IM vaccination, hamsters were anesthetized, fur over the rear thigh removed, and the Tropis placed over the semitendinosus and biceps femoris muscles for activation of the device.

### Hantavirus challenge

Female Syrian hamsters (Envigo) were anesthetized by inhalation of vaporized isoflurane using an IMPAC 6 veterinary anesthesia machine. Once anesthetized, hamsters were injected with 200 PFU of ANDV, SNV, SEOV, HTNV, DOBV, or 1,000 PFU PUUV diluted in phosphate-buffered saline (PBS). IM (caudal thigh) injections consisted of 0.2 ml delivered using a 1-ml syringe with a 25-gauge, 5/8-inch needle.

### Pseudovirion neutralization assay (PsVNA)

The PsVNA using a non-replicating VSVΔG-luciferase pseudovirion system was performed as previously described [15].

### N-ELISA

The ELISA using to detect N-specific antibodies (N-ELISA) was described previously [13, 24]. The endpoint titer was determined as the highest dilution that had an optical density (OD) greater than the mean OD for serum samples from negative control wells plus 3 SDs. The PUUV N antigen was used to detect SNV and PUUV N-specific antibodies [20] and SEOV N antigen was used to detect SEOV, HTNV, and DOBV N-specific antibodies [13].

### ISH

A probe targeting the M segment messenger RNA was synthesized and purchased from Advanced Cell Diagnostics (ACD, Newark, CA). After deparaffinization with xylene, a series of ethanol washes, and peroxidase blocking, sections were heated in antigen retrieval buffer and then digested by proteinase. Sections were exposed to the probe and incubated at 40 °C in a hybridization oven for 2 h. After rinsing, ISH signal was amplified using kit-provided Pre-amplifier and Amplifier conjugated to alkaline phosphatase, and incubated with Fast Red substrate solution for 10 min at room temperature. Sections were then stained with hematoxylin, air-dried, and mounted.

### IFA

Formalin-fixed paraffin embedded (FFPE) tissue sections were deparaffinized using xylene and a series of ethanol washes. After 0.1% Sudan black B (Sigma) treatment to eliminate the autofluorescence background, the sections were heated in Tris-EDTA buffer (10mM Tris Base, 1mM EDTA solution, 0.05% Tween 20, pH 9.0) for 15 min to reverse formaldehyde crosslinks. After rinses with PBS (pH 7.4), the sections were blocked with PBS containing 5% normal goat serum overnight at 4 °C. The sections were incubated with either rabbit anti-SEOV (1:200), mouse anti-vimentin (1:500), mouse anti-desmin (1:100), rat anti-CD45 (1:100), rat anti-F4/80 (1:100), or mouse anti-alpha smooth muscle actin (1:500) for 2 hr at room temperature. After rinses with PBS, sections were incubated with either secondary goat anti-rabbit Alexa Fluor 488 (green, 1:500), goat anti-rat Alexa Fluor 568 (red, 1:500), or goat anti-mouse Alexa Fluor 647 (magenta, 1:500), for 1 hr at room temperature. Sections were cover slipped using the Vectashield mounting medium with DAPI (Vector Laboratories). Images were captured on a Zeiss LSM 880 confocal system and processed using ImageJ software.

### Expression and Potency IFA

Plasmid DNA was combined at a 3:1 ratio of FuGENE^®^ 6 Transfection Reagent (Promega) to DNA and incubated 20 min at RT. Two-fold serial dilutions were made and added to 293T cells seeded on black, clear bottom plates then incubated 18 to 24 hours at 37°C. Cells were fixed with 1:1 methanol/acetone for 10 min at −20°C and blocked using fish gelatin (Rockland Labs) for 30 min at 37°C. Plates were incubated with a rabbit anti-ANDV (from a rabbit vaccinated with pWRG/AND-M by muscle electroporation, 1:500) for 1 hour at 37°C, washed 3 times with PBS-T, and incubated with a goat anti-rabbit Alexa Fluor 488 (1:400) for 30 min at 37°C. Washed plates were read immediately using an EVOS FL fluorescent microscope.

### Ethics

Animal research was conducted under an IACUC approved protocol at USAMRIID (USDA Registration Number 51-F-00211728 & OLAW Assurance Number A3473-01) in compliance with the Animal Welfare Act and other federal statutes and regulations relating to animals and experiments involving animals. The facility where this research was conducted is fully accredited by the Association for Assessment and Accreditation of Laboratory Animal Care, International and adheres to principles stated in the Guide for the Care and Use of Laboratory Animals, National Research Council, 2011.

### Statistical analysis

Comparison of titers was done using Student’s *t* test (two-tailed). *P* values of less than 0.05 were considered significant. Survival analyses were done using log-rank test. Analyses were conducted using GraphPad Prism (version 8).

## Results

### Hantavirus DNA vaccines administered by needle and syringe

Hantavirus DNA vaccines evaluated in Syrian hamsters using needle and syringe have been poorly immunogenic [13, 25]. For example, hamsters vaccinated with a SEOV DNA vaccine with needle and syringe elicited some neutralizing antibody; however, the response was dramatically lower than the same vaccine delivered by gene gun [13]. To determine if this poor response could be overcome by plasmid construct optimization, an experiment was conducted to evaluate the immunogenicity of hantavirus DNA vaccines that had been optimized for codon-usage and mRNA stability. Groups of 7 or 8 hamsters were vaccinated with 200 ug of either the non-optimized ANDV DNA vaccine, optimized ANDV DNA vaccine, optimized SNV DNA vaccine, or optimized SEOV DNA vaccine delivered at 4-week intervals for a total of 4 vaccinations (**Fig. 1A**). Sera collected throughout the vaccination series were analyzed for neutralizing antibodies using a homotypic PsVNA. Following 4 vaccinations, all hamsters vaccinated with the non-optimized ANDV DNA vaccine, the optimized ANDV DNA vaccine, and the optimized SNV DNA vaccine were negative by PsVNA (**Fig. 1B**). When subjected to the homologous virus challenge there was not a statistically significant difference between vaccinated and unvaccinated animals in survival of ANDV-infected hamsters (**Fig. 1C**) or N-ELISA titers of SNV-infected hamsters (**Fig. 1D**). Of the hamsters vaccinated with the optimized SEOV DNA vaccine, 3/7 hamsters had positive PsVNA50 titers following the 4^th^ vaccination (**Fig. 1B**). After challenge with SEOV, 5/7 hamsters were protected from infection as determined by N-ELISA on Day 28 (**Fig. 1E**). The poor immune response after needle/syringe delivery, even for optimized vaccines, confirmed our earlier findings [13] that another method of delivering DNA was needed for use in small animals.

**Fig. 1.**
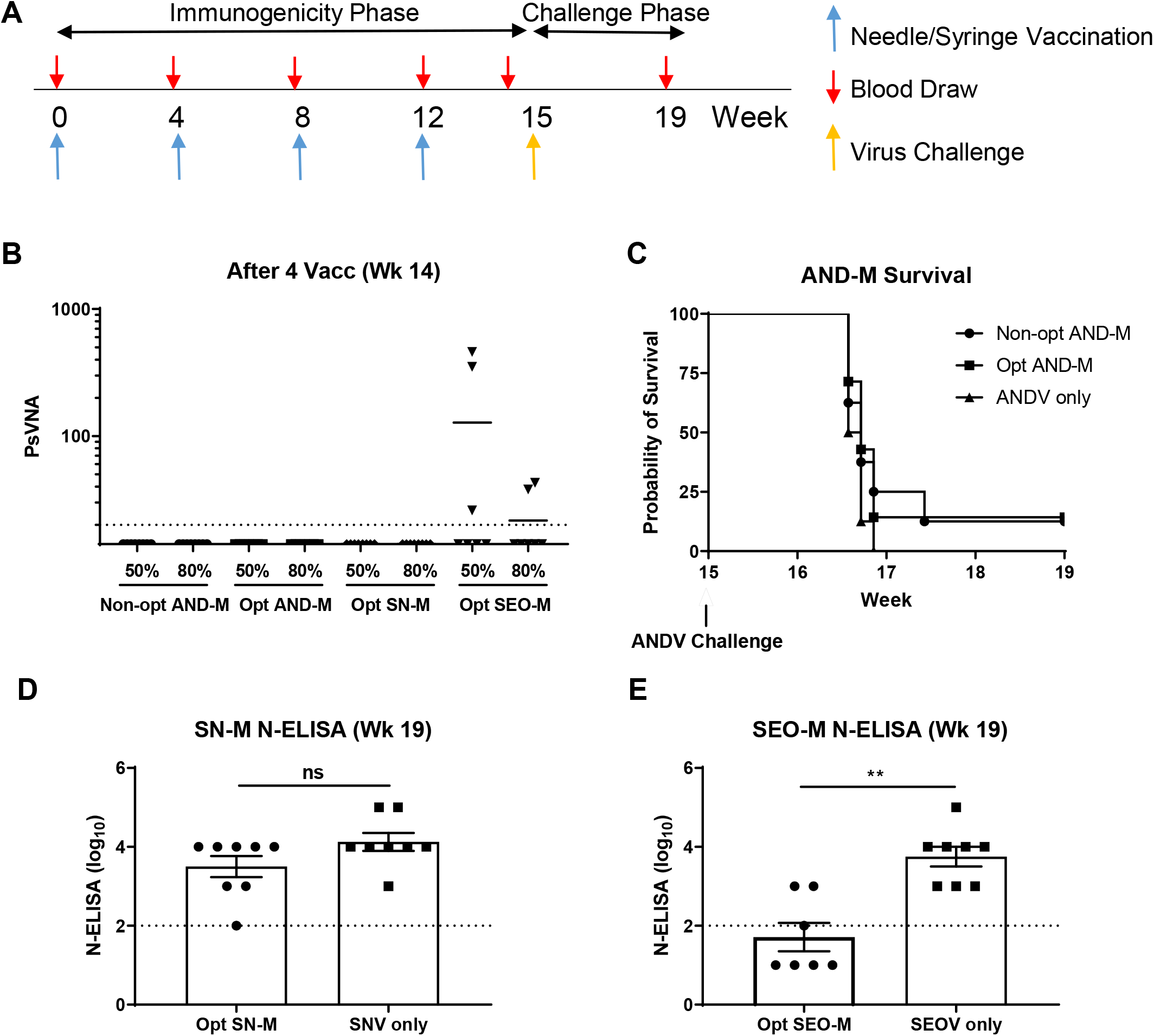
Low immunogenicity of hantavirus DNA vaccines administered by needle/syringe. A) Groups of 8 hamsters were vaccinated with 200ug of either the non-optimized ANDV DNA vaccine, optimized ANDV DNA vaccine, optimized SNV DNA vaccine, or optimized SEOV DNA vaccine delivered 4 times at 4-week intervals using needle/syringe. **B)** Hantavirus strain-specific PsVNA titers were determined after 4 vaccinations. The PsVNA limit of detection was a titer of 20 (dashed line). **C)** Hamsters vaccinated with non-optimized and optimized ANDV vaccines were subsequently challenged with 200 PFU ANDV. Survival of all animals was monitored for 4 weeks postchallenge. **D)** Serum from SNV DNA vaccinated hamsters 4 weeks following a 200 PFU SNV challenge was analyzed by N-ELISA and titers plotted. **E)** Serum from SEOV DNA vaccinated hamsters 4 weeks following a 200 PFU SEOV challenge was analyzed by N-ELISA and titers plotted. The N-ELISA limit of detect was a titer of a 2 log10 (dashed line). **, *P* < 0.01; ns, not significant.

### SEOV DNA vaccine administered by PharmaJet^®^ Tropis

The same hantavirus DNA vaccines that were poorly immunogenic in hamsters vaccinated with needle and syringe were highly immunogenic when delivered by jet injection in larger species such as rabbits, geese, nonhuman primates [15, 26]. We were interested in adapting a jet injection technology for use in smaller animals such as hamsters. The approach we took was to evaluate the PharmaJet^®^ Tropis human intradermal delivery device in hamsters. Two techniques of using Tropis to deliver the vaccine were tested. The first technique involved pinching shaved abdominal skin such that the skin was stretched over the gloved forefinger. The Tropis device was then pressed against the skin over the forefinger and activated (**Fig 2A**). The second technique involved pressing the Tropis device into the shaved skin overlying the semitendinosus and biceps femoris muscles (**Fig 2B**). To determine grossly where the inoculum was delivered, PBS mixed with a contrasting dye, and delivered using the pinched-skin technique or the over-the-muscle technique. Visualization of the dye indicated that the pinched-skin technique resulted in delivery of the dye to the dermis and SC spaces (data not shown). The over-the-muscle technique resulted in intradermal, SC and successful IM injection (data not shown). The pinched-skin technique will henceforth be referred to as SC route and the over-the-muscle technique will be referred to as IM route.

**Fig. 2.**
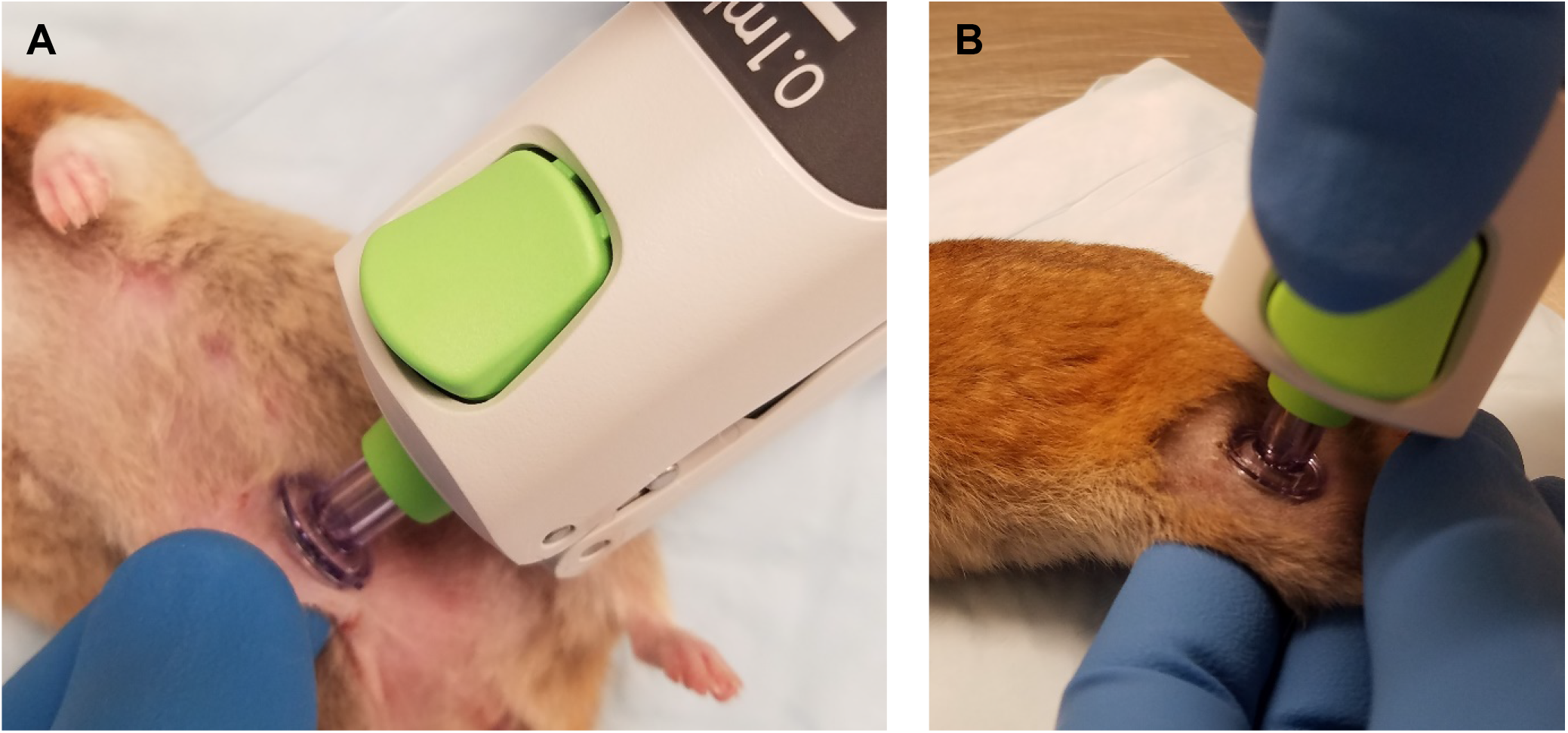
Tropis SC and IM techniques. **A)** SC technique image depicting the pinching of abdominal skin for vaccine administration using Tropis. **B)** IM technique image depicting placement of Tropis for IM vaccine administration.

The optimized SEOV DNA vaccine was selected for the initial PharmaJet^®^ Tropis experiment to evaluate the two different techniques for delivering a DNA vaccine. The SEOV DNA vaccine was used so a comparison could be made between Tropis and the low (but detectable) responses elicited when vaccine was delivered by needle/syringe. Groups of 8 hamsters were vaccinated 3 times with the optimized SEOV DNA vaccine by either the SC or IM routes (**Fig. 3A**). Serum collected following the 2^nd^ and 3^rd^ vaccination and prior to virus challenge were analyzed by PsVNA (**Fig. 3B**). Unexpectedly, high neutralizing antibody titers were obtained following both the 2^nd^ (geometric mean titers [GMT] of 4,858 PsVNA50 for IM and 4,065 PsVNA50 for SC) and 3^rd^ (GMT 11,915 PsVNA50 for IM and 6,704 PsVNA50 for SC) vaccinations. There was not a statistically significant difference comparing the two routes of vaccination; and there was only a statistically significant difference in titers between the 2^nd^ and 3^rd^ vaccination by the IM route. Increased variability in titers was observed with SC vaccination, thereby highlighting the IM route as the more consistent method of vaccination for future experiments. All hamsters vaccinated with the optimized SEOV DNA vaccine, regardless of route, were protected from infection when challenged with SEOV (**Fig. 3C**).

**Fig. 3.**
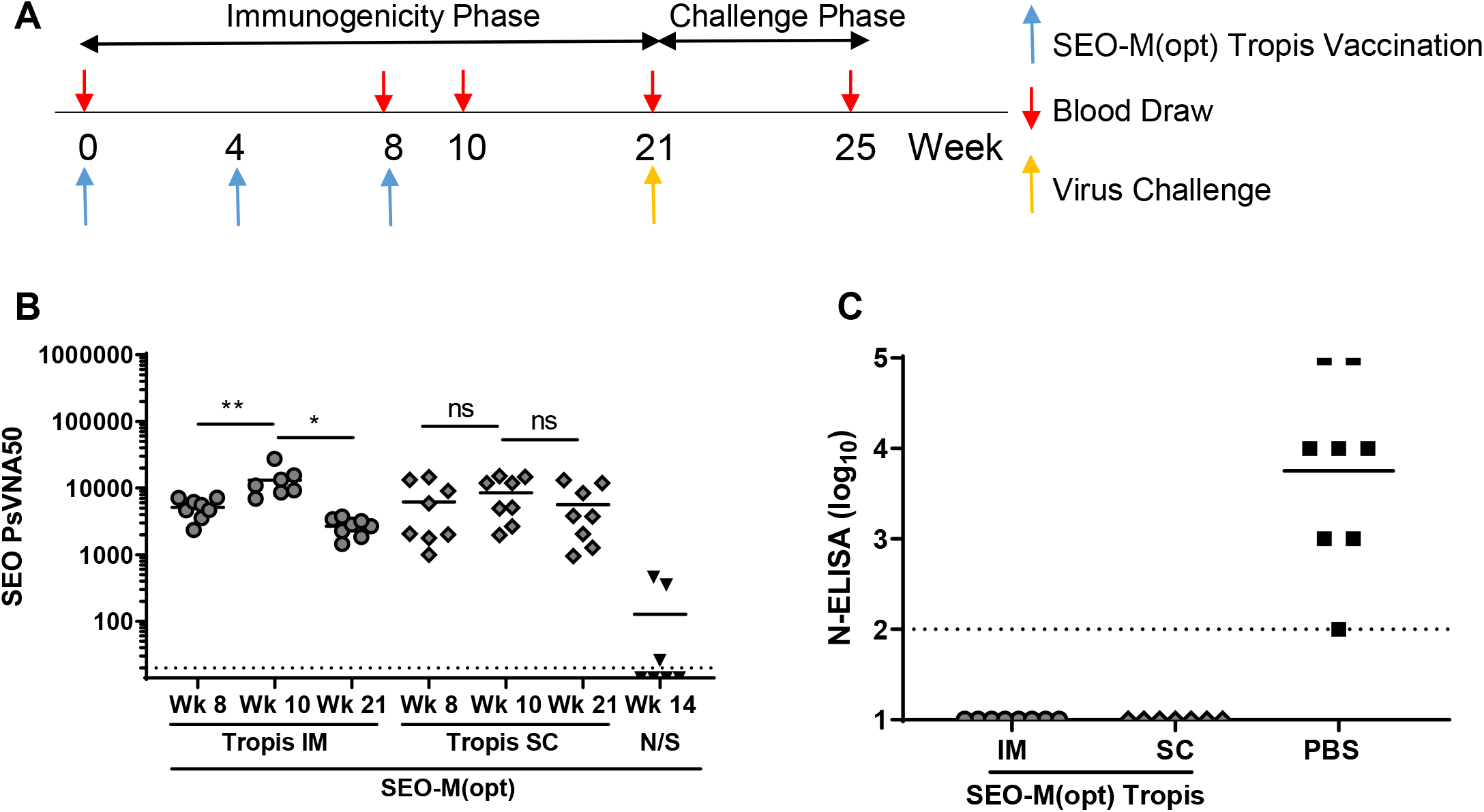
SEOV DNA vaccine immunogenicity in hamsters using PharmaJet Tropis. **A)** Groups of 8 hamsters were vaccinated with 200ug of the optimized SEOV DNA vaccine 3 times at 4-week intervals using the PharmaJet Tropis at either the IM or SC site. **B)** SEOV PsVNA titers were determined on weeks 10 and 21 post-vaccination. Week 14 SEOV DNA vaccine by needle/syringe (N/S) data from **Fig.** 1 are illustrated for comparison. The PsVNA limit of detection was a titer of 20 (dashed line). **C)** Serum from SEOV DNA vaccinated hamsters 4 weeks following a 200 PFU SEOV challenge was analyzed by N-ELISA and titers plotted. The N-ELISA limit of detect was a titer of a 2 log10 (dashed line). *, *P* < 0.05; **, *P* < 0.01; ns, not significant.

Tissues from the injection site and inguinal lymph nodes were collected 2 days post-vaccination and analyzed for pathological findings and by ISH. A single animal injected with optimized SEOV DNA vaccine via needle/syringe at the SC site had a significant finding by H&E (**Fig. 4A**). A dermal focal area, positive using a probe targeting the M segment RNA sequence, admixed with lymphocytes, macrophages, and rare eosinophils were observed (**Fig. 4B**). Using the PharmaJet^®^ Tropis device, positive staining was only observed in epidermis and subadjacent dermis in hamsters vaccinated by either the IM or SC technique (**Fig. 4C, D**). All inguinal lymph nodes were negative; however, in select animals, connective tissue and fibroblasts surrounding the lymph node were positive (**Table 1**). Staining was scored by intensity and show slightly higher staining comparing the PharmaJet^®^ Tropis vaccination device over needle/syringe delivery.

**Table 1.** ISH comparing pWRG/SEO-M(opt2) vaccine delivery systems.

| Vaccine | Delivery System | Vaccination Route | ISH Vaccination Site Score<sup>a</sup> | ISH Inguinal Lymph Node Score<sup>b</sup> |
| --- | --- | --- | --- | --- |
| pWRG/SEO-M(opt2) | PharmaJet® Tropis | IM | +, ++, ++ | Neg, +++, Neg |
| pWRG/SEO-M(opt2) | PharmaJet® Tropis | SC | +, ++, +++ | +, Neg, + |
| pWRG/SEO-M(opt2) | Needle/Syringe | IM | +, +, + | All Negative |
| pWRG/SEO-M(opt2) | Needle/Syringe | SC | +++, ++, + | All Negative |
| PBS | PharmaJet® Tropis or Needle/Syringe | IM/SC | All Negative | All Negative |

<sup>a</sup> Positive cell types include epidermis and dermis.
<sup>b</sup> Positive cell types include connective tissue and fibroblasts, all lymph nodes negative.

**Fig. 4.**
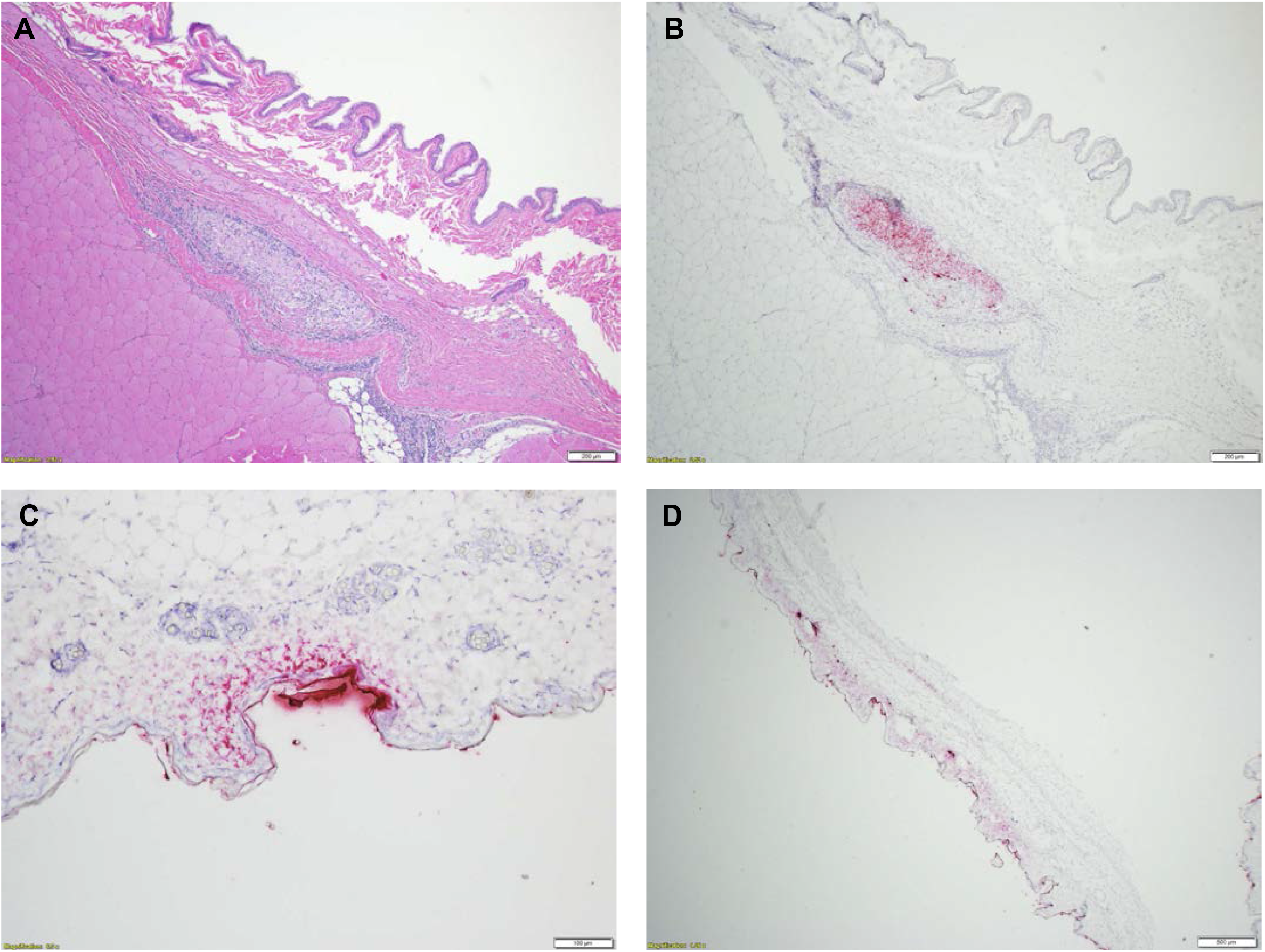
Optimized SEOV DNA vaccine RNA In Situ Hybridization at Vaccination Sites. Hamsters were vaccinated with 200ug of the optimized SEOV DNA vaccine using either the PharmaJet Tropis or Needle/Syringe. **A)** H&E stain (4x) and **B)** ISH (4x) of hamster vaccinated using Needle/Syringe demonstrating positive staining in the dermis. **C)** Hamster vaccinated using the PharmaJet Tropis by the IM route demonstrating positive staining from the epidermis to the subadjacent dermis (10x). **D)** Hamster vaccinated using the PharmaJet Tropis by the SC route demonstrating diffuse, positive staining from epidermis to dermis (2x).

Antibodies targeting specific cell types (*i.e*. muscle cells, leukocytes, macrophages, fibroblasts) were used to determine the cells expressing SEOV antigen. By immunofluorescence assay, it was determined that cell types expressing SEOV antigen were vimentin-positive cells (**Fig. 5A**), further supporting the ISH results. Co-staining was not observed in muscle cells (desmin-positive cells, **Fig. 5B**), leukocytes (CD45-positive cells (**Fig. 5B**), macrophages (F4/80-positive cells, **Fig. 5C**), or myofibroblasts (αSMA-positive cells, **Fig. 5D**).

**Fig. 5.**
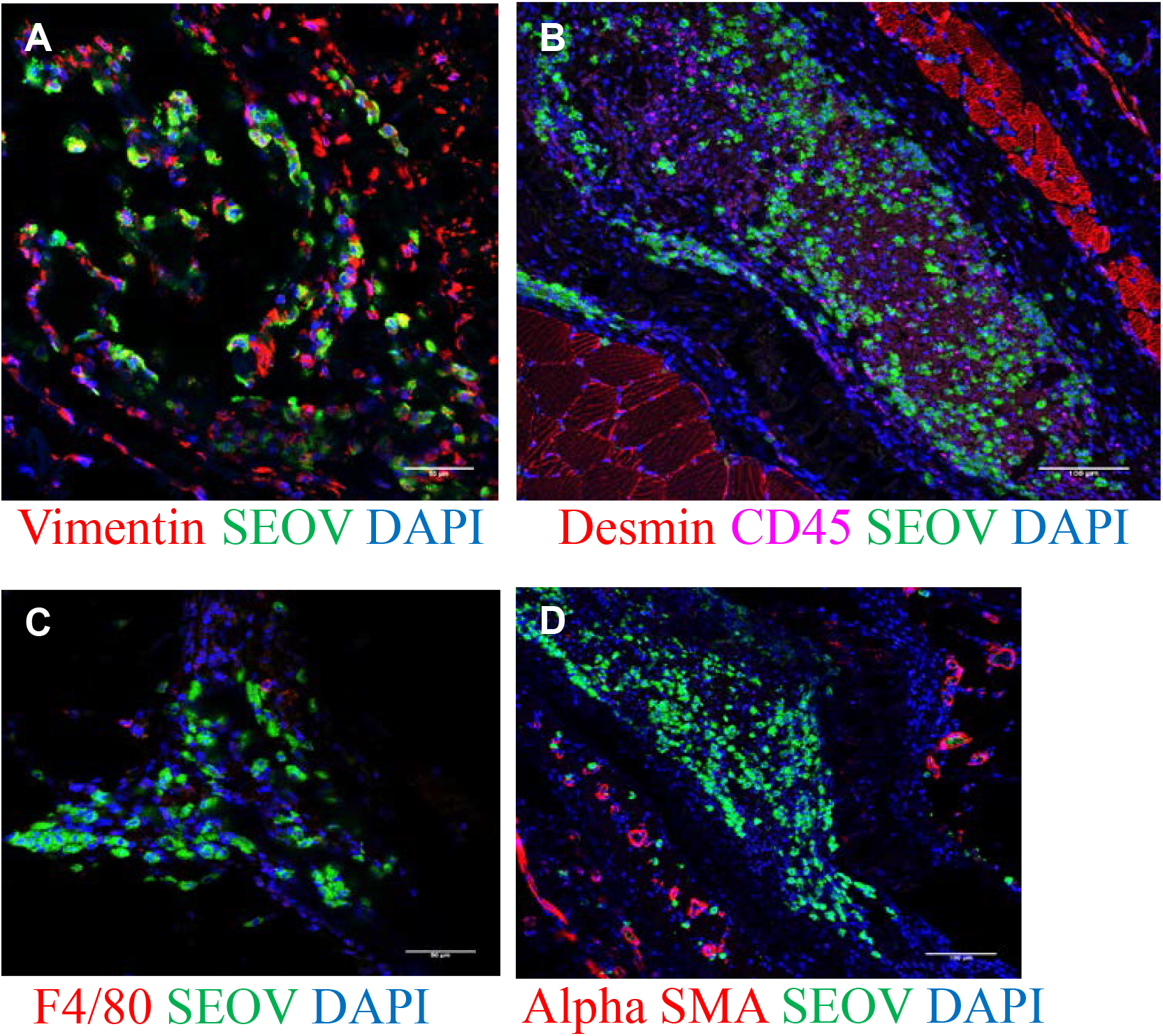
SEOV antigen colocalizes with vimentin-positive cells. Hamsters were vaccinated with 200ug of the optimized SEOV DNA vaccine using the PharmaJet Tropis at the SC site. FFPE tissue sections were stained with **A)** anti-vimentin and anti-SEOV antibodies, **B)** anti-desmin, anti-CD45 and anti-SEOV antibodies, **C)** anti-F4/80 and anti-SEOV antibodies, and **D)** anti-SMA and and anti-SEOV antibodies. Scale bars (**A-D**), 50 μm.

### Effect of gene optimization on hantavirus DNA vaccine immunogenicity

Comparisons of three non-optimized and optimized hantavirus DNA plasmids were analyzed in hamsters vaccinated using the PharmaJet^®^ Tropis vaccination device. These included pWRG/SEO-M, pWRG/AND-M(1.1), and pWRG/HTN-M(x) (non-optimized) versus pWRG/SEO-M(opt2), pWRG/AND-M(opt2), and pWRG/HTN-M(co) (optimized), respectively. Surprisingly, there was only a statistically significant difference following the 3^rd^ vaccination when comparing the non-optimized and the optimized SEOV DNA vaccines (PsVNA50 GMT 1539 vs. 11915, respectively, p=0.0222) (**Fig. 6A, B**). Optimization of the ANDV DNA vaccine did result in an increased number of animals with positive PsVNA50 titers following the third vaccination (1/7 vs. 9/16 animals); however, the increase in PsVNA GMT did not reach statistical significance (PsVNA50 GMT <20 vs. 62, respectively, p=0.1018) (**Fig. 6D, E**). There was a 4.7-fold increase in PsVNA50 titers observed in hamsters after the 3^rd^ vaccination comparing the non-optimized and optimized HTNV DNA vaccines, yet this increase did not reach statistical significance (PsVNA50 GMT 1038 vs. 4981, respectively, p=0.2345) (**Fig. 6G, H**).

**Fig. 6.**
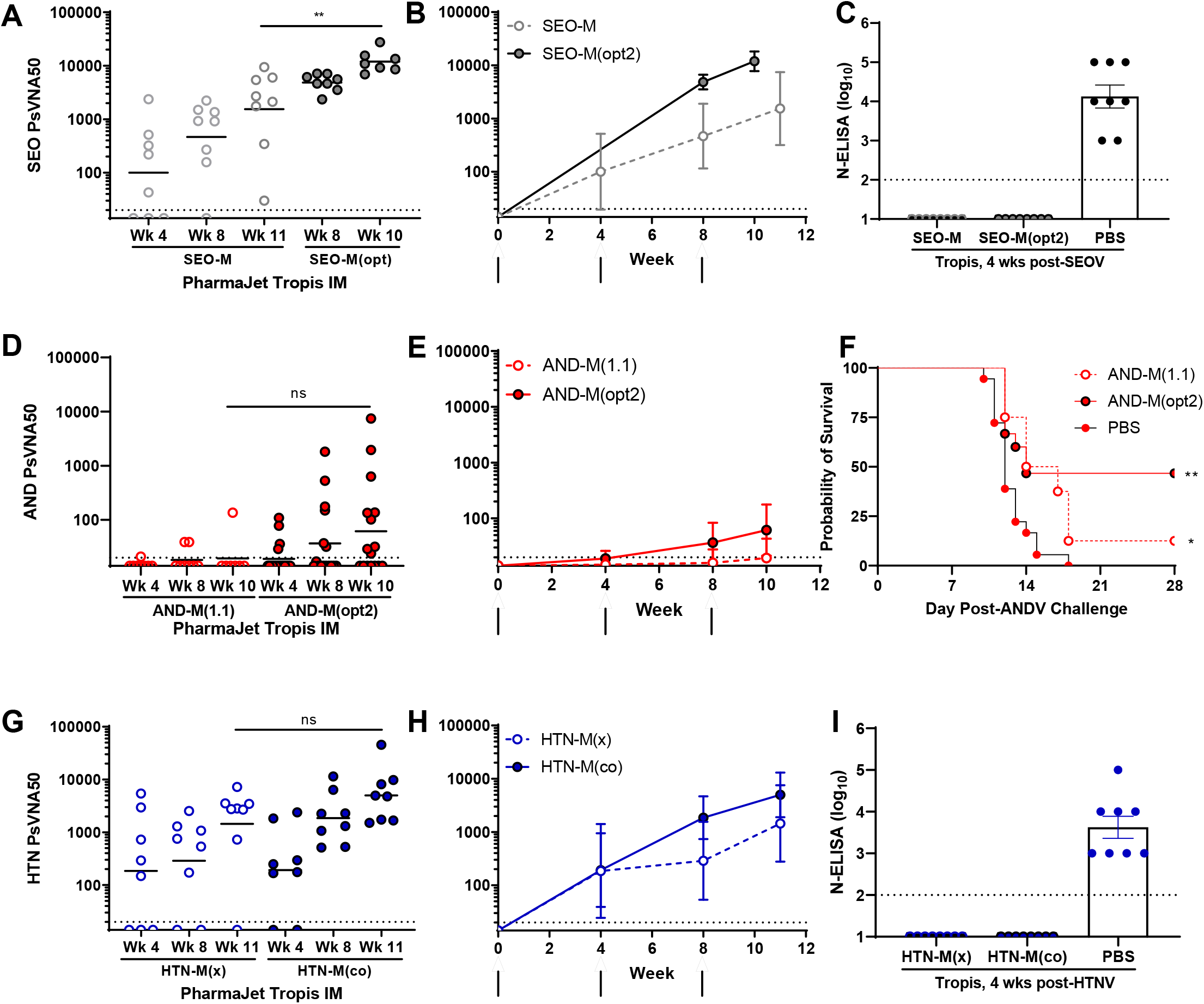
Optimization does not necessarily result in increased immunogenicity. Groups of 8 hamsters were vaccinated with 200ug of either the non-optimized and optimized SEOV DNA vaccines **(A, B, C),** non-optimized and optimized ANDV DNA vaccines, **(D, E, F)** or non-optimized and optimized HTNV DNA vaccines **(G, H, I)** 3 times at 4-week intervals using the PharmaJet Tropis at the IM site. Individual PsVNA titers are plotted **(A**, **D**, **G)** and geomean titers over the vaccination course (**B**, **E, H)** and **F)**. N-ELISA titers are reported for SEOV and HTNV infection models **(C, I)**. Survival data is displayed for the ANDV/hamster disease model **(F)**. The PsVNA limit of detection was a titer of 20 (dashed line). The N-ELISA limit of detect was a titer of a 2 log10 (dashed line). **, *P* < 0.01; ns, not significant.

Following the vaccination series, hamsters were challenged with 200 PFU of the homotypic virus. All hamsters vaccinated with either the non-optimized or optimized SEOV or HTNV DNA vaccines were protected from infection as measured by N-ELISA (**Fig. 6C, I**). Of the hamsters challenged with ANDV, only 1/8 animals vaccinated with the non-optimized ANDV DNA vaccine survived (**Fig. 6F**). There was, however, a statistically significant delay in death (p=0.0137). Of the hamsters vaccinated with the optimized ANDV DNA vaccine, 7/15 (a single animal from this group was found dead prior to ANDV challenge) animals survived to the end of the study (p=0.0014). This is the first demonstration of protection of hamsters by active vaccination with any ANDV DNA vaccine.

### Immunogenicity of hantavirus DNA vaccines administered using PharmaJet^®^ Tropis

Titers and seropositivity of hantavirus DNA vaccines administered using PharmaJet^®^ Tropis was compared to historical vaccination using gene gun (**Table 2**). Comparable titers (within a 2-fold difference) were observed between non-optimized SEOV, ANDV, and HTNV DNA vaccines by the two vaccination methods. In contrast, the optimized hantavirus DNA vaccines, targeting SNV and PUUV had an approximately 8- and 63-fold increase in immunogenicity (p=0.0013 and p<0.0001, respectively). All hamsters vaccinated with the optimized SNV, PUUV, and DOBV DNA vaccines and subsequently challenged with the homologous virus were completely protected from infection (**Fig. 7A-F**).

**Table 2.**
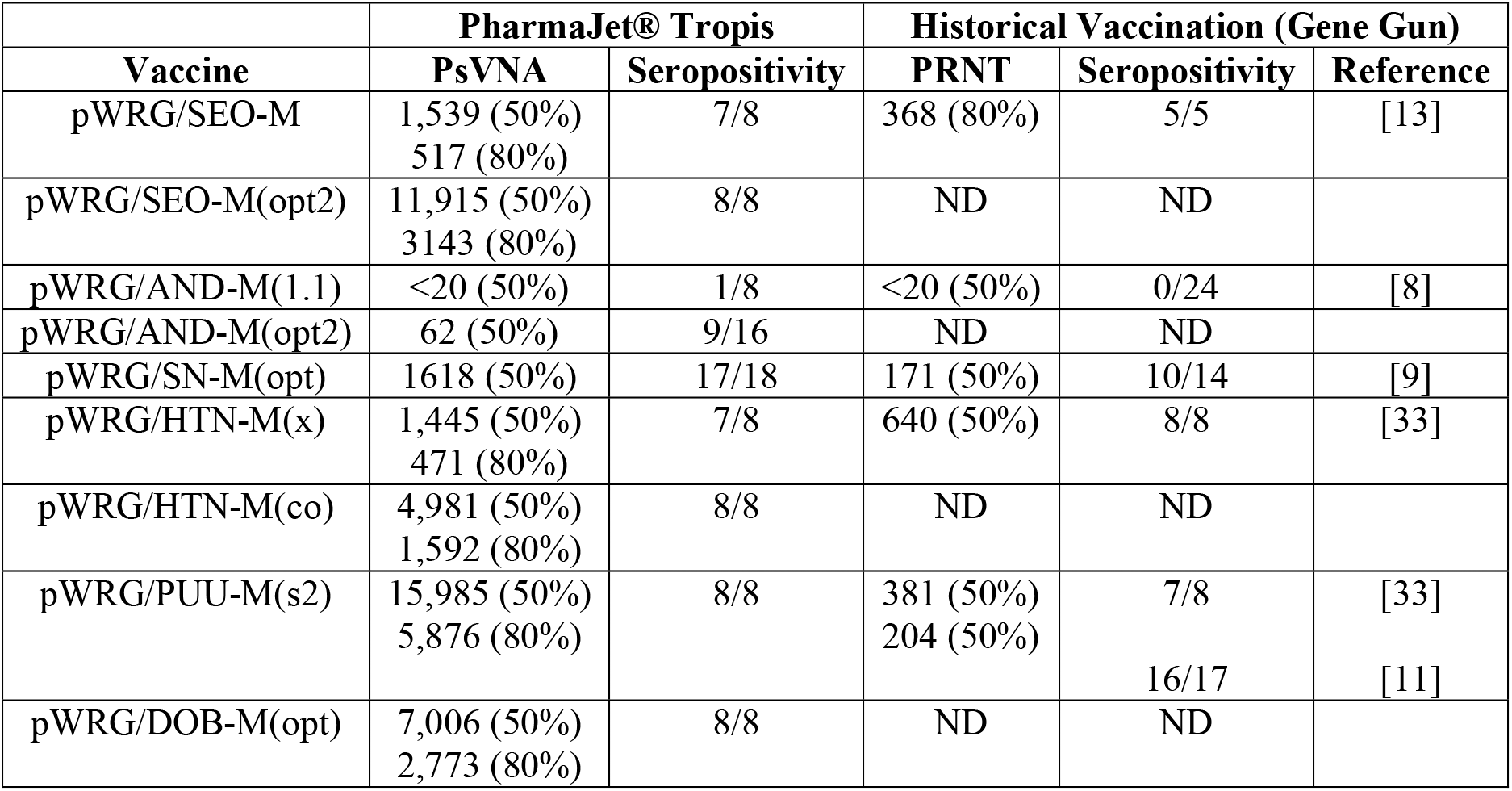
Hantavirus DNA vaccine titers using PharmaJet^®^ Tropis and Gene Gun.

**Fig. 7.**
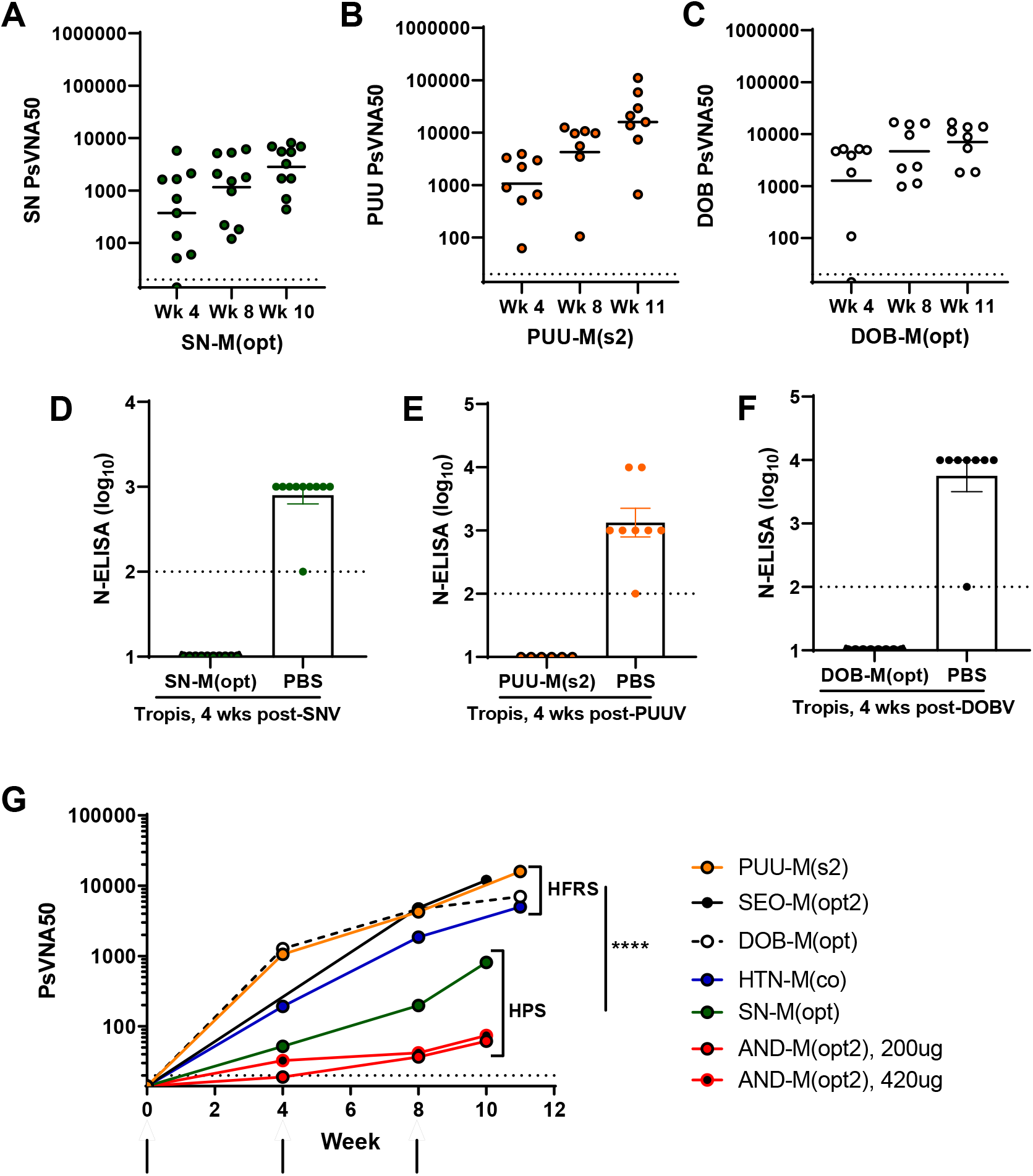
Immunogenicity and protection of optimized hantavirus DNA vaccines. Groups of 8-10 hamsters were vaccinated with 200ug of the indicated hantavirus DNA vaccine using PharmaJet Tropis on weeks 0, 4 and 8. Individual PsVNA titers are plotted (**A-C**). The PsVNA limit of detection was a titer of 20 (dashed line). Serum collected 28 days following either SNV, PUUV or DOBV challenge was analyzed by N-ELISA (**D-F**). The N-ELISA limit of detect was a titer of a 2 log10 (dashed line). **G)** Groups of 8 hamsters were vaccinated with either 200ug or 420ug of the ANDV DNA vaccine using PharmaJet Tropis at the IM site on weeks 0, 4 and 8 (black arrows). Serum was analyzed using the homotypic PsVNA. The PsVNA limit of detection was a titer of 20 (dashed line). ****, *P* < 0.0001.

The immunogenicity of all optimized hantavirus DNA vaccines administered using the PharmaJet^®^ Tropis were compared in hamsters (**Fig. 7G**). There is a stark contrast between the immunogenicity of hantavirus DNA vaccines targeting HFRS-causing hantaviruses when compared to hantavirus DNA vaccines targeting HPS-causing hantaviruses (Week 10/11, PsVNA50 GMT 8948 vs. 187, respectively, p<0.0001).

We evaluated the non-optimized and optimized ANDV DNA vaccines in an *in vitro* potency assay typically used for evaluating hantavirus DNA vaccines for stability in clinical trials. The optimized ANDV DNA vaccine had an approximate 2-fold increase in potency when compared to the non-optimized ANDV DNA vaccine (data not shown). However, given the reduced titers of the optimized ANDV DNA vaccine administered by PharmaJet^®^ Tropis in hamsters, a second experiment was conducted to evaluate an increased dosage based on the concentration of the vaccine and the volume restrictions of the vaccine delivery device (**Fig. 7G**). After 3 vaccinations, there was not a statistically significant difference between hamsters administered 420 ug or 200 ug optimized ANDV DNA vaccine by PharmaJet^®^ Tropis. An equivalent number of animals per vaccination dosage (6/10 animals from each group) survived ANDV exposure.

## Discussion

Despite the technical challenges of testing hantavirus DNA vaccines in small animals, several hantavirus DNA vaccines are currently in clinical trials (reviewed in [18]). The evolution of these hantavirus DNA vaccines is synonymous with DNA vaccine development, in general. The overall goal of enhancing antigen expression through delivery efficiency, codon optimization, and formulation/adjuvant has advanced these vaccines forward to the current state of animal/human trials.

The PharmaJet^®^ Stratis and Tropis are FDA 510(k)-cleared devices and have been used or are currently in-use for several ongoing DNA vaccine clinical trials. In a Phase I clinical trial for two experimental Zika virus DNA vaccines, vaccination with the Stratis device resulted in statistically significant increased immunogenicity over needle/syringe delivery [27]. Similarly, delivery of ANDV and SNV DNA vaccines with the PharmaJet^®^ Stratis device elicited statistically significant increases in antibody titers when compared to IM needle/syringe delivery in rabbits and nonhuman primates [15]. Clinical trials evaluating the safety of HTNV, PUUV, and ANDV DNA vaccines delivered by PharmaJet^®^ Stratis are ongoing [18].

The Tropis device is a needle-free jet injection system that we have shown can be adapted for use in small animal studies providing a cost benefit to the initial development of vaccines. In this study, however, we have observed that Tropis, advertised as an intradermal injection on humans, penetrates through the dermis and deeper when used on a Syrian hamster recapitulating an IM or SC injection. This in fact more closely resembled a Stratis injection of nonhuman primates or humans. The PharmaJet^®^ Stratis device can either administer vaccine as an IM or SC injection. The exact cell type that takes up injected plasmid DNA after jet injection is not fully understood. Using a series of antibodies that cross-react with hamster antigens, we were able to identify the epidermis, dermis, and fibroblast cells that either take up the DNA or express the antigen.

We evaluated the effect of gene optimization to enhance immunogenicity of hantavirus DNA vaccines in hamsters. Gene optimization capitalizes on different organisms’ bias toward codon usage for the same amino acid. This strategy was used to enhance immunogenicity of DNA vaccines targeting avian influenza virus [28, 29] and HIV-1 [30, 31]. We initially pursued optimization as a potential strategy to overcome hantavirus vaccine interference [32, 33]. The use of the PharmaJet^®^ Tropis allows for a comparison of non-optimized and optimized hantavirus DNA vaccines. Others have noted that codon optimization does not always correlate with vaccine efficacy [34]. In this study, the only statistically significant difference observed was between the non-optimized SEOV DNA vaccine and optimized SEOV DNA vaccine (**Fig. 6A**). Previously, during the development of the PUUV DNA vaccine, several amino acid differences were noted between the initial first-generation vaccine and published sequences [11]. Vaccination of hamster and nonhuman primates with the first-generation PUUV DNA vaccine and a synthetic, optimized PUUV DNA vaccine resulted in none or very limited immunogenicity. This was overcome by replacing 5 consensus residues from the first-generation vaccine into the optimized vaccine resulting in enhanced immunogenicity. Taken together, our experience with optimized vaccines shows that they have the potential to increase the immunogenicity of the vaccine, but this is not always the case.

The disparity of titers measured between hamsters vaccinated with HFRS and HPS vaccines may be species-specific. Previous studies using rabbits and nonhuman primates do not replicate the significant difference observed when hamsters were vaccinated with hantavirus DNA vaccines using the PharmaJet^®^ Tropis [9, 15]. While titers from HPS DNA vaccines (SNV and ANDV DNA vaccines) are significantly lower than HFRS DNA vaccines, all hamsters vaccinated with the SNV DNA vaccine were protected from infection. Hamsters vaccinated with the optimized ANDV DNA vaccine with titers >60 PsVNA50 at the time of challenge were protected from lethal ANDV challenge. This is a significant increase from ANDV DNA vaccination using other vaccine delivery systems.

## Conclusions

In conclusion, we have identified a novel vaccine delivery system approach to test hantavirus DNA vaccines in hamsters. The PharmaJet^®^ Tropis device, delivering a volume of 0.1 mL, makes this delivery system feasible for small animal studies. Using this device, we have demonstrated immunogenicity and protective efficacy of both HFRS and HPS DNA vaccines in hamsters. This work would not have been possible with needle and syringe. The ease of use of the PharmaJet^®^ Tropis system will enable future experiments to determine why HPS DNA vaccines have reduced immunogenicity when compared to HFRS DNA vaccines, and to test formulation/adjuvanting methods in a small animal species. This approach can be used to further evaluate hantavirus DNA vaccine formulation or adjuvants and can be used for other DNA vaccines that utilize a hamster model, such as vaccines targeting COVID-19.

## Acknowledgements

We thank the USAMRIID Comparative Medicine Division and Pathology Division for technical assistance. The authors also thank David Fetterer for statistical analysis. This work was supported by the Military Infectious Disease Research Program (MIDRP), Program Area T. Opinions, interpretations, conclusions, and recommendations are ours and are not necessarily endorsed by the U.S. Army or the Department of Defense. No competing interests declared. The funders had no role in the study design, data collection and analysis, decision to publish or preparation of the manuscript.

## Author Contributions

Conceptualization: Rebecca L. Brocato, Jay W. Hooper Investigation: Rebecca L. Brocato, Steven A. Kwilas, Matthew D. Joselyn, Simon Long, Casey C. Perley, Lucia M. Principe, Brandon Somerville, Melanie V. Cohen, Jay W. Hooper Methodology: Rebecca L. Brocato, Jay W. Hooper Analysis: Rebecca L. Brocato, Steven A. Kwilas, Simon Long, Kevin Zeng, Jay W. Hooper Writing (original draft): Rebecca L. Brocato Writing (reviewing and editing): Steven A. Kwilas, Matthew D. Joselyn, Simon Long, Kevin Zeng, Casey C. Perley, Lucia M. Principe, Brandon Somerville, Melanie V. Cohen, Jay W. Hooper

